# Co-inhibition of BCL-XL and MCL-1 with BCL-2 selective inhibitors A1331852 and S63845 enhances cytotoxicity of cervical cancer cell lines

**DOI:** 10.1101/824649

**Authors:** Siti Fairus Abdul Rahman, Kalaivani Muniandy, Yong Kit Soo, Elvin Yu Huai Tiew, Ke Xin Tan, Timothy E. Bates, Nethia Mohana-Kumaran

## Abstract

A combination of the BCL-2 inhibitors ABT-263 and A-1210477 inhibited cell proliferation in the HeLa, C33A, SiHa and CaSki human cervical cancer cell lines. Drug sensitivity was initially tested using 2-dimensional (2D) cell culture models. As ABT-263 binds to both BCL-2 and BCL-XL at high affinity, it was unclear whether the synergism of the drug combination was driven either by singly inhibiting BCL-2 or BCL-XL, or inhibition of both. Therefore, we used the BCL-2 selective inhibitor ABT-199 and the BCL-XL selective inhibitor A1331852 to resolved the individual antitumor activities of ABT-263 into BCL-2 and BCL-XL dependent mechanisms. A-1210477 was substituted with the orally bioavailable S63845. The SiHa, C33A and CaSki cell lines were resistant to single agent treatment of all three drugs, suggesting that none of these anti-apoptotic proteins singly mediate survival of the cells. HeLa cells were resistant to single agent treatment of ABT-199 and A1331852 but were sensitive to S63845 indicating that they depend on MCL-1 for survival. Co-inhibition of BCL-XL and MCL-1 with A1331852 and S63845 significantly inhibited cell proliferation of all four cell lines. Similar data were obtained with 3-dimensional spheroid cell culture models generated from two cervical cancer cell lines *in vitro*. Treatment with a combination of A1331852 and S63845 resulted in inhibition of growth and invasion of the 3D spheroids. Co-inhibition of BCL-2 and MCL-1 with ABT-199 and S63845, also inhibited cell proliferation of all cancer cell lines, except SiHa. However, the effect of the combination was not as pronounced as combination of A1331852 and S63845. Collectively, our data demonstrate that the combination of MCL-1-selective inhibitors with either selective inhibitors of either BCL-XL or BCL-2 may be potentially useful as treatment strategies for the management of cervical cancer.

## 1. Introduction

The BCL-2 family proteins are crucial regulators of the intrinsic apoptosis pathway. These proteins can be divided into pro-apoptotic and anti-apoptotic proteins and have one to four BCL-2 homology motifs (BH1-BH4). The anti-apoptotic multidomain (BH1-BH4) members namely BCL-2, BCL-XL, BCL-w, BFL-1/A1 and MCL-1 function to counteract the pore-forming activity of the pro-apoptotic multidomain proteins (BH1-BH4), BAX and BAK which permeabilize the mitochondria outer membrane. Following various stress signals, the BH3-only proteins either neutralize the anti-apoptotic proteins or directly activate effector proteins BAX and BAK which will eventually lead to apoptosis in cells [1, 2].

One strategy that cancer cells employ to evade apoptosis, triggered by oncogenesis or drug treatment is via overexpressing the BCL-2 anti-apoptotic proteins [3]. Hence, treatment that is effective in activating pro-death signaling either by upregulating pro-apoptotic protein BIM or effector proteins BAX or BAX are inefficient as cancer cells can survive the cytotoxic insult by sequestering pro-apoptotic proteins with anti-apoptotic proteins [4]. Cellular anti-apoptotic mechanisms can also be suppressed by BCL-2 selective inhibitors [4], which mimic the action of certain BH3-only proteins. For example, ABT-263 (Navitoclax) mimics the BH3-only protein BAD which selectively inhibits BCL-2, BCL-XL and BCL-w [5]. ABT-263 has also demonstrated anti-tumor activity in lymphoid malignancies in clinical studies, but induced dose-dependent thrombocytopenia as a consequence of inhibiting BCL-XL [6, 7]. This toxicity prompted the development of the BCL-2 selective inhibitor ABT-199/venetoclax [8]. Venetoclax was approved by FDA for the treatment of chronic lymphocytic leukemia (CLL) [9] but has shown activity in other cancers such as acute myeloid leukemia (AML) [10] and T-cell acute lymphoblastic leukemia (T-ALL) in combination with the MCL-1 selective inhibitor S63845 [11]. In order to determine the contribution of BCL-XL for survival of cancer cells, a number of specific BCL-XL inhibitors such as WEHI-539 [12], A1331852 and A1155463 [13] have been developed.

In this present study, ABT-199 and A1331852 [13] were used experimentally to investigate the contributions of BCL-2 and BCL-XL in mediating cervical cancer cell survival. In order to study the role of MCL-1 for cell survival, S63845, a small molecule inhibitor of MCL-1 was employed. S63845 was reported to demonstrate higher affinity towards MCL-1 (K_i_ < 1.2nM) compared to A-1210477 (K_i_ = 28 nM). In addition, S63845 was 1000-fold more potent in killing (MCL-1 dependent) H929 cells compared to A-1210477 [14], and its use therefore would be more appropriate in helping delineate its role in cervical cancer cell survival.

Four cervical cancer cell lines C33A, SiHa, HeLa and CaSki were subjected to single agent treatment of ABT-199, A1331852 and S3845. The cells were also tested with combinations of A1331852/S63845 and ABT-199/S63845 in monolayer (2D) culture and in 3-dimensional (3D) spheroids, which provide a microenvironment closer to tumours *in vivo* [15].

## 2. Material and Methods

### 2.1 Drugs and Cell Lines

ABT-199, A1331852 and S63845 (MedChemExpress, NJ, USA) were dissolved in dimethyl sulfoxide (DMSO) at a stock concentration of 10 mM. All four cell lines were purchased from the American Type Culture Collection (Manassas, VA, USA), and maintained in culture as described previously [16].

### 2.2 Drug sensitivity assay

Drug sensitivity assays were performed as described previously [17]. Cells were first subjected to ABT-199, A1331852 and S63845 treatment alone or in combination for 72 hours. Sensitivity of cells to drug combinations was measured by testing a fixed concentration of S63845 with increasing concentrations of either A1331852 or ABT-199. Cell proliferation was quantified by fluorescence using SYBR Green as described previously [16]. All drug sensitivity assays were conducted four times (n = 4) and average IC_**50**_ values were calculated from the experimental data.

### 2.3 Three-dimensional spheroids

Approximately 5000 cells (2.5 × 10^4^ cells/ml) cells were seeded in an Ultra-Low Attachment (ULA) 96-well U bottom-plate (Corning, NY, USA). Plates containing the cells were centrifuged at 1200 rpm for 2 minutes. Plates were incubated at 37°C, 95% O_**2**_, 5% CO_2_ for 72 hours. After 72 hours, 3D spheroids were embedded into collagen mix [18]. Spheroids were treated with A1331852, ABT-199 and S63845, alone and in combination for 72 hours. Spheroid growth and invasion were photographed every 24 hours using a Nikon C2+ inverted confocal microscope. Upon termination of the assay, live-dead staining of spheroids was conducted as described in [19]. Images were taken using a Nikon-300 inverted fluorescence microscope.

## 3. Results

### 3.1 Selective BCL-2 inhibitors resolve the individual contributions of anti-apoptotic proteins BCL-2, BCL-XL and MCL-1 for cervical cancer cell lines survival

HeLa cells were resistant to single agent treatment with A1331852 (Fig. 1a & Table S1) and ABT-199 (Fig. 1b & Table S1) but sensitive to single agent treatment with S63845 (Fig. 1c & Table S1). C33A (Fig. 1a-c & Table S1), and SiHa (Fig. 1a-c & Table S1) cells were resistant to single agent treatment with all three BCL-2 selective inhibitors. CaSki cells were slightly sensitive to A1331852 (Fig. 1a & Table S1), but were resistant to single agent ABT-199 (Fig. 1b & Table S1) and S63845 (Fig. 1c & Table S1). Although slightly sensitive to A1331852, it appears that co-inhibition of other anti-apoptotic proteins are necessary for more effective killing of the CaSki cells.

**Fig. 1:**
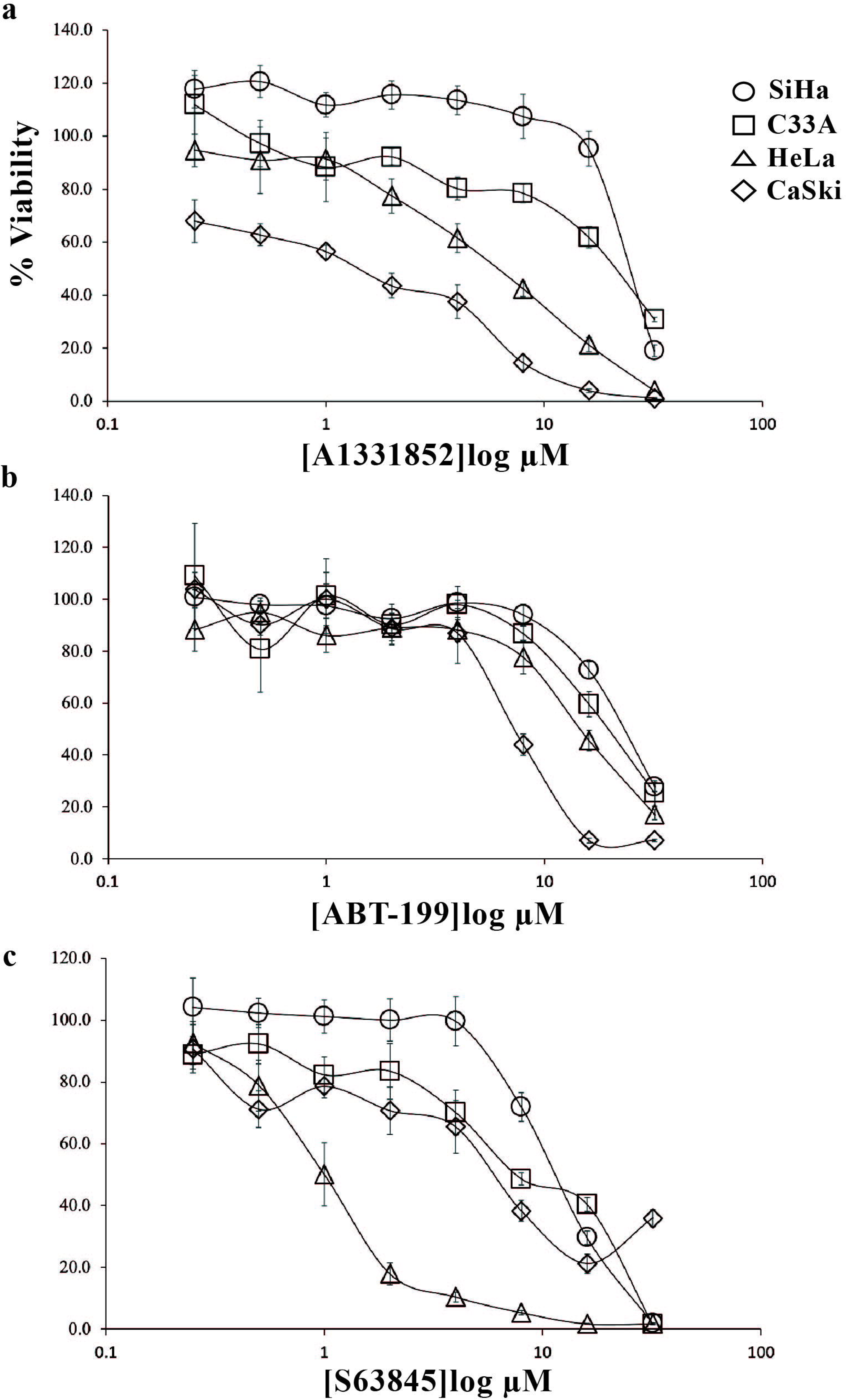
Sensitivity of the cervical cancer cell lines to single agent treatment of ABT-199, A1331852 and S63845. **(a)** HeLa, C33A and SiHa were resistant to single agent treatment of A1331852. CaSki cells were slightly sensitive to A1331852; **(b)** All four cell lines were resistant to single agent treatment of ABT-199. **(c)** Except for HeLa, all other cervical cancer cell lines were insensitive to single agent treatment of S63845. Points represent mean ± SEM of four experiments.

Collectively, these data suggest that insensitivity of HeLa cells to single agent treatment of ABT-199 and A1331852 shows that they depend on MCL-1 for survival, as the cells were susceptible to single agent treatment of S63845. Insensitivity of the other cell lines to all three BCL-2 selective drugs used as monotherapy suggest that the cells are resistant to apoptosis due to the need to target multiple pro-survival proteins rather than just one. These data also suggest that other death mechanisms and pathways may be responsible for apoptotic death mechanisms in these cells. For example, it has been demonstrated that there are non-caspase dependent cell death mechanisms that are dependent on the cathepsins [20].

### 3.2 Substantial inhibition of cell proliferation driven by co-inhibition of BCL-XL and MCL-1

As HeLa cells were sensitive to single agent S63845 (Fig. 1c & Table S1), we tested the sensitivity of HeLa to fixed doses of S6835 (doses below 1 µM) with increasing concentrations of either A1331852 or ABT-199.

HeLa cells were treated with either a fixed dose of 0.25 µM or 0.5 µM S63845 and increasing concentrations of A1331852 (0 - 32 µM). At a concentration of 0.25 µM S63845, the dose-response curve substantially moved to the left (Fig. 2a). 0.25 µM S63845 sensitized HeLa cells to A1331852 by 44-fold (Table S2). Similar data were obtained when the concentration of S63845 was increased to 0.5 µM (Fig. 2a & Table S2). In C33A cells, the presence of 0.5 µM S63845, shifted the dose-dependent curve substantially to the left (Fig. 2B & Table S2) and sensitized the cells to A1331852 close to 100-fold (Table S2). Addition of 1 µM and 2 µM S63845 (Fig. 2b & Table S2) resulted in similar data. Comparably, in the presence of 0.5 µM S63845 (Fig. 2c), SiHa cells were sensitized to A1331852 by 100-fold (Table S2). Similar data were obtained in SiHa cells when the concentration of S63845 was increased to 1 µM and 2 µM (Fig. 2c & Table S2). In CaSki cells, combination with S63845 sensitised the cells to A1331852 for all concentrations tested (Fig. 2d & Table S2) indicating that co-inhibition with MCL-1, enhances cell killing compared to inhibition of BCL-XL alone. The CI values obtained for combination of A1331852 and S63845 exhibited synergism at several concentrations for all four cervical cancer cell lines (Table S3).

**Fig. 2:**
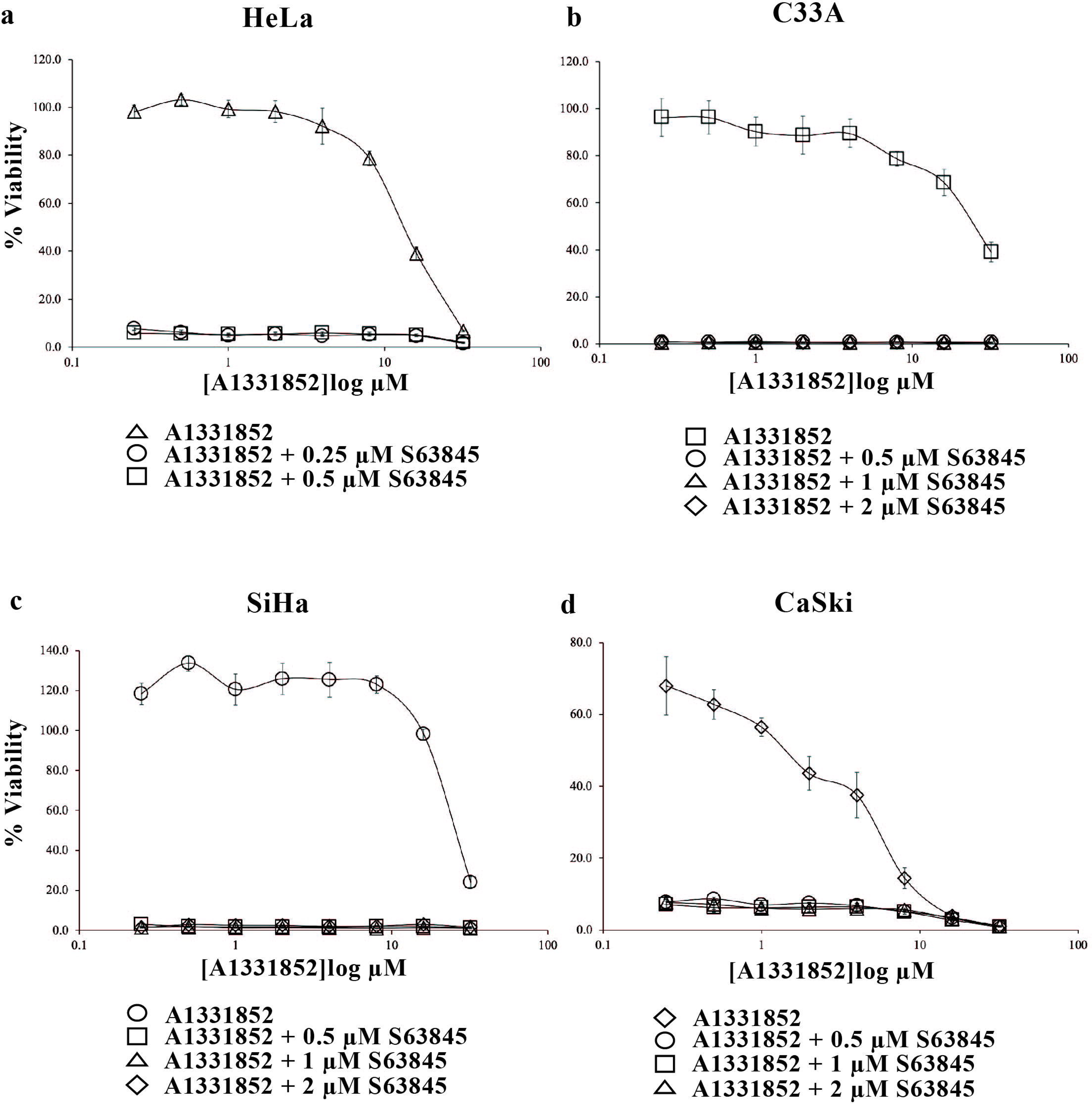
Co-inhibition of BCL-XL and MCL-1 using BCL-2 selective inhibitors A1331852 and S63845. Cervical cancer cell lines **(a)** HeLa; **(b)** C33A; **(c)** SiHa and **(d)** CaSki cells were treated with increasing concentrations of A1331852 (0-32 µM) in the presence and absence of S63845. Points represent mean ± SEM of four experiments.

### 3.3 S63845 sensitized 3-dimensional (3D) spheroids generated from cervical cancer cell lines to A1331852 but not to ABT-199

S63845 and A1331852 used as single agents had less effect on the growth and invasion of the spheroids except at 1 µM of S63845 (Fig. 3 – see yellow box) and 1 µM of A1331852 (Fig. 3 – see green box), there was a noticeable decrease in viability in the periphery of the spheroids. When combined, in the presence of 1 µM of S63845, there was obvious sensitization of the spheroids to A1331852. This manifested as reduced spheroid growth and invasion (Fig. 3 – see the column in red). Taken together, the synergistic effect of the drug combination on growth and invasion of the spheroids was similar to the cytotoxicity curves obtained for the 2D cultures (Fig. 2a-c). Similar data were obtained when the drug combination was tested on 3D spheroids generated from SiHa cells. S63845 at 2 µM was able to sensitize the spheroids to A1331852, reflected in dose-dependent inhibition of spheroid growth and invasion. Similarly, A1331852 at 2 µM was able to sensitize the spheroids to S63845 (Fig. S1). Taken together, the effect of combination of A1331852/S63845 observed in the spheroid model was consistent with the monolayer culture data, suggesting that this drug combination may be effective *in vivo*.

**Fig. 3:**
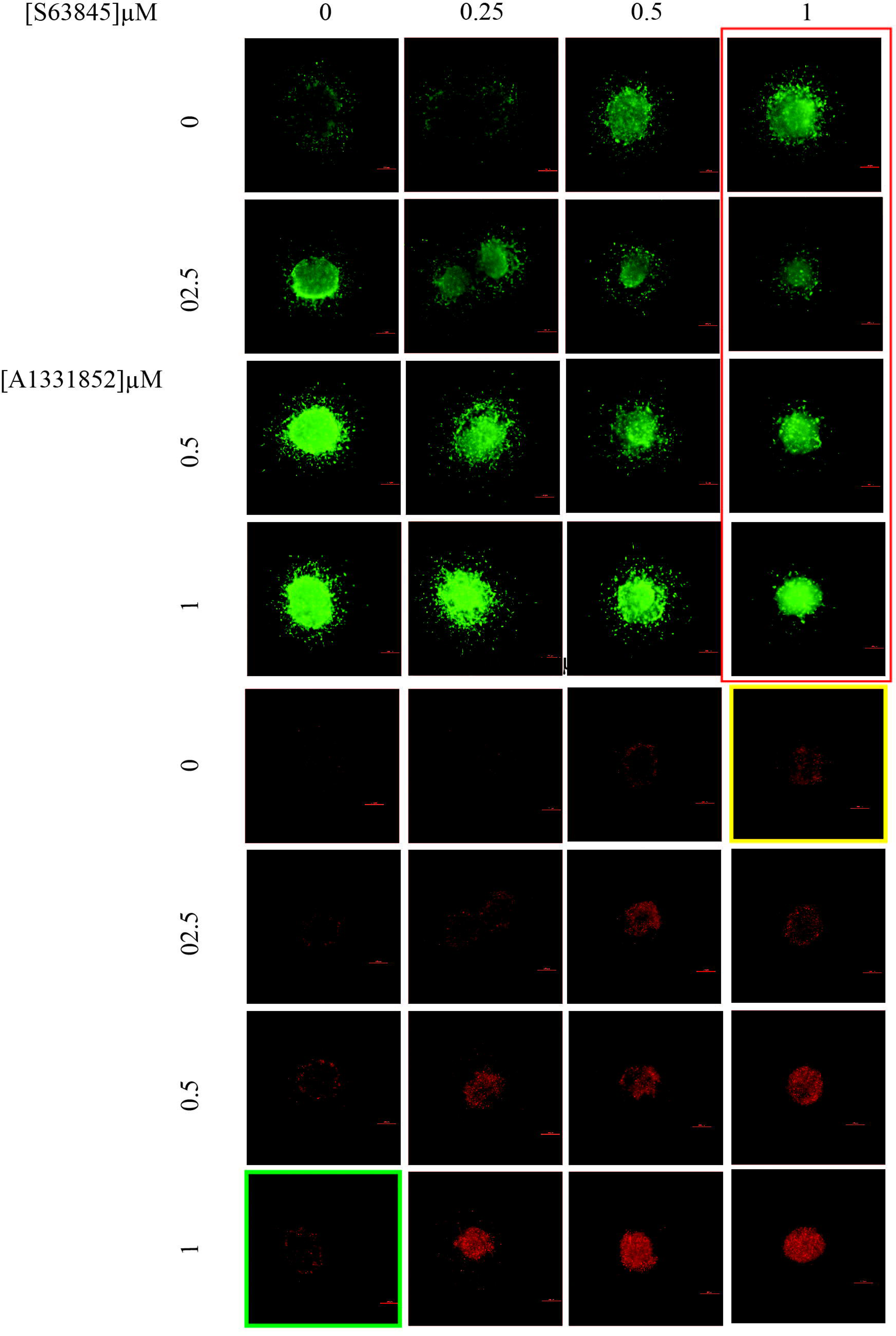
The effect of combination of S63845 and A1331852 on the growth and invasion of 3D HeLa spheroids over three days. The spheroids were treated with single agents S63845 and A1331852 and combination of both over three days at the indicated concentrations. Cell viability was determined using the live/dead assay (Viable cells: stained green by Calcein-AM; Dead cells: stained red by Ethidium-homodimer I). Size bar: 200 µm.

S63845 only modestly sensitised HeLa cells to ABT-199 in monolayer culture (Fig. 2a). The combination (ABT-199/S63845) however, had minimal effect on the growth and invasion of the 3D HeLa spheroids even at the highest combination concentration used, indicating higher combination concentrations may be required to inhibit growth and invasion of the spheroids (Fig. S2).

### 3.4 Cervical cancer cell lines were sensitive to co-inhibition of BCL-2 and MCL-1

In HeLa cells, 0.25 µM S63845 shifted the dose-response curve to the left (Fig. 4a) sensitizing the cells to ABT-199 by 6-fold (Table S4). An increase in concentration of S63845 to 0.5 µM, resulted in a significant shift of the dose-response curve to the left (Fig. 4a) and the cells were sensitized to ABT-199 by 13-fold (Table S4).

**Fig. 4:**
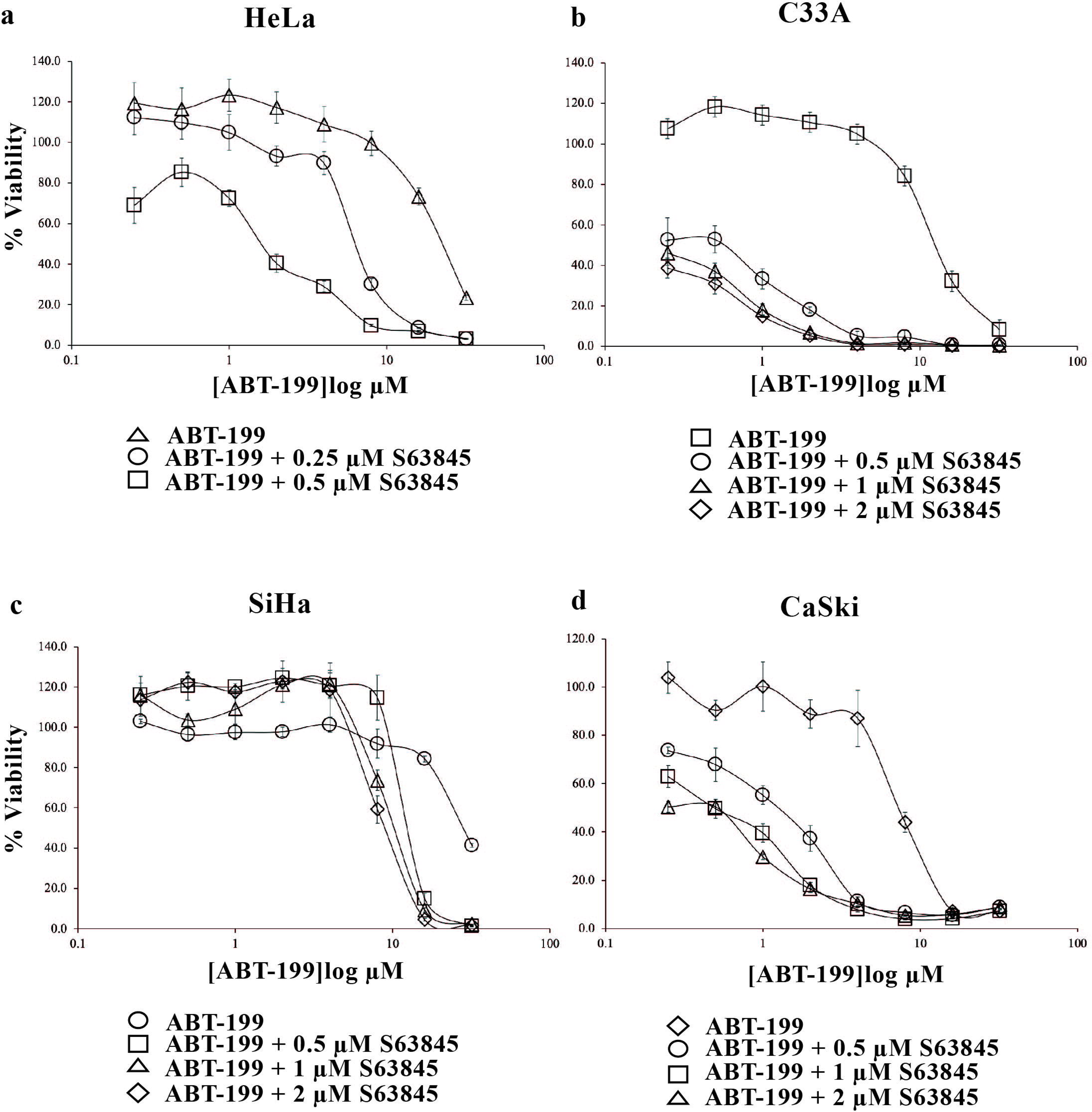
Co-inhibition of BCL-2 and MCL-1 using BCL-2 selective inhibitors ABT-199 and S63845. Cervical cancer cell lines **(a)** HeLa; **(b)** C33A; **(c)** SiHa and **(d)** CaSki cells were treated with increasing concentrations of ABT-199 (0-32 µM) in the presence and absence of S63845. Points represent mean ± SEM of four experiments.

The drug interaction analyses demonstrated that combination of ABT-199 with 0.25 µM of S63845 could not be determined. At the concentrations tested, the poor efficacy of the combination treatment, meant that we were unable to conduct drug interaction analyses (Table S5). Concentrations of ABT-199 > 1 µM combined with 0.25 µM S63845 were antagonistic (Table S5). The combination of S63845 only resulted in synergism at 0.5 µM S63845 combined with concentrations of ABT-199 > 1 µM (Table S5). The words “antagonism”, and “synergism” refer to the overall effect on cell proliferation and are not in any way meant to infer the properties of a classical pharmacological ligand that is an antagonist (in relation for example to a cell surface receptor and agonists/antagonists) pharmacological sense.

At 0.5 µM S63845, C33A cells were sensitized to ABT-199 by 22-fold (Fig. 4b & Table S4). The sensitization increased to > 40-fold at a concentration of 1 µM S63845 and 2 µM of S63845 (Fig. 4b & Table S4) and drug interaction analyses demonstrated strong synergism at multiple doses of S63845 and ABT-199 (Table S5).

In SiHa cells, S63845 at 0.5 µM (Fig. 4c & Table S4) and 1 µM (Fig. 4c) only sensitized SiHa cells to ABT-199 by 2-fold (Table S4). This sensitization only increased to 3-fold (Table S4) when the concentration of S63845 was increased to 2 µM (Fig. 4c).

Combination with 0.5 µM of S63845 modestly sensitised the CaSki cells to ABT-199 by 6-fold (Fig. 4d & Table S4). The fold-sensitisation increased (14 - fold), when concentration of S63845 was increased to 1 µM and 2 µM (Fig. 4d & Table S4). Drug interaction analyses indicated that the drug combinations demonstrated strong synergism at several concentrations of S63845 and ABT-199 (Table S5). Collectively, the findings demonstrate that inhibition of either BCL-2 or BCL-XL alone is not adequate to kill the CaSki cells. Co-inhibition of MCL-1 with either BCL-XL or BCL-2 appears to be essential to kill the cells.

These data demonstrate that there was a greater response to co-inhibition of MCL-1 and BCL-XL. Cells responded to combination of S63845 and A1331852 more rapidly at low concentrations. In contrast, the response to co-inhibition of MCL-1 and BCL-2 was variable suggesting that other cell death mechanisms that do not rely on these proteins may be involved.

## 4. Discussion

Our data suggest that in all cell lines tested co-inhibition of MCL-1 is important and necessary to induce cell death, as none of the cell lines responded to ABT-263 used singly. However, ABT-263 is reported to cause thrombocytopenia due to BCL-XL inhibition [6, 7]. Hence, it is important to investigate whether selective inhibition of BCL-XL or BCL-2 would minimize toxicity and serve as a substitute for ABT-263. Hence, we employed the BCL-2 selective inhibitors ABT-199, A1331852 and S63845 to define the contributions of these anti-apoptotic proteins in maintaining survival of the cervical cancer cells.

All four cervical cancer cell lines tested were resistant to single agent treatment of ABT-199 and A1331852. None of the cell lines, except HeLa responded to S63845, when used singly, indicating that they were not solely MCL-1-dependent. However, although HeLa cells responded to single agent treatment of S63845, treatment with a combination of ABT-199 or A1331852 with concentrations of S63845 of < 1 µM resulted in synergy, indicating that inhibition of either BCL-XL or BCL-2 is still required to achieve cell killing at lower doses of S63845. These data demonstrate that survival of the cervical cancer cell lines is maintained by more than one anti-apoptotic protein and selectively inhibiting them in combination kills the cells more effectively.

A number of studies have also shown that survival of cancer cells is dependent on the expression of several different anti-apoptotic proteins. For example, chronic lymphocytic leukemia (CLL) cells are killed when either BCL-2 and BCL-XL or BCL-2 and MCL-1 [21]. Acute myeloid leukemia (AML) cells developed resistance to inhibition of BCL-2 by upregulating BCL-XL and MCL-1. Therefore, inhibiting both BCL-XL and MCL-1 resensitized AML cells to ABT-199, which inhibits BCL-2 [22]. Furthermore, co-inhibition of MCL-1 and BCL-2 killed T-ALL cells *in vitro* and *in vivo* [11]. These present data show that all four cervical cancer were sensitive to combinations of A1331852 and S63845 at lower combination concentrations, indicating that they depend on both BCL-XL and MCL-1 for survival, and co-inhibition of these molecules are sufficient to cause cell death. Moreover, our data show that BCL-XL is the key target of ABT-263, for inducing the synergy observed previously with A-1210477.

The sensitization obtained in the monolayer culture was analogous to the data obtained with the 3D spheroid studies. The 3D HeLa spheroids were sensitized to A1331852 by S63845 but sensitization was only obvious following treatment with 1 µM of S63845, indicating that higher concentrations of S63845 are required to sensitize spheroids to A1331852 compared to concentration of S63845 required to see the same sensitization effect in monolayer culture. One explanation for the need of higher drug combination concentrations in the spheroids, could be attributed to the 3D orientation of the tumor cells which is likely to limit diffusion of drugs to the cells in the center of the spheroid.

C33A cells were sensitive to combinations of ABT-199 and S63845. Given that the C33A cells also responded effectively to a combination of A1331852 and S63845, it appears that co-inhibition of either BCL-2 or BCL-XL with MCL-1 is sufficient to trigger cell death in C33A cells. SiHa cells responded poorly to combination of ABT-199 and S63845 but the cells were sensitive to combination of A1331852 and S63845. Therefore, SiHa cells may be dependent on BCL-XL and MCL-1 for survival rather than BCL-2. Therefore, it is possible that that co-inhibition of BCL-2 and MCL-1 may have led to overexpression of BCL-XL as a compensatory survival adaptation which has been reported in other cancer cell lines. CLL cells developed resistance to ABT-737 (which selectively inhibits BCL-2 and BCL-XL) treatment due to concurrent upregulation of BCL-XL and BFL-1/A1 [23] and upregulation of MCL-1 and BFL-1/A1 resulted in acquired resistance in a number of cancer cells to ABT-737 [19, 24, 25].

All four cell lines were more responsive to lower doses of combination of S63845 and A1331852 compared to combination of S63845 and ABT-199, indicating that BCL-XL and MCL-1 are better targets for inducing cervical cancer cell line killing. Other studies have also demonstrated that inhibition of BCL-XL rather than BCL-2 has resulted in sensitization of solid tumor cancer cell lines to other drugs. For example, the BCL-XL inhibitor WEHI-539 but not BCL-2 inhibitors sensitized osteosarcoma cell lines to doxorubicin [26]. Breast cancer, non-small cell lung cancer ovarian cancer cell lines were sensitized to docetaxel by ABT-263 and BCL-XL selective inhibitors but not to BCL-2 inhibitors [13]. Chondrosarcoma cell lines were reported to be sensitized to doxorubicin or cisplatin by BCL-XL inhibitors and not BCL-2 inhibitors both *in vitro* and *in vivo* [27]. More recently, drug combinations targeting BCL-XL and MCL-1, and to a lesser extent BCL-2 were reported to synergistically kill melanoma cells in 2D and 3D cell culture models [28]. Collectively, solid tumors may be more susceptible to inhibition of BCL-XL and MCL-1. However, co-targeting BCL-XL and MCL-1 may pose an issue in the clinic, as inhibition of BCL-XL may result in thrombocytopenia [6, 7]. At present neither A1331852 nor S63845 are useful in the clinic, due to toxicity issues and co-targeting of BCL-XL and MCL-1 can cause fatal hepatotoxicity [29]. However, our present data suggest that selective, less toxic BCL-XL inhibitors may be useful in combination with conventional chemotherapy and/or the use of selective pro-apoptotic agents that directly activate type 2 mitochondrial pathways [2]. Another strategy would be to co-inhibit BCL-2 and MCL-1.

Testing the drug combinations used here in rodent models are necessary for determining safety and efficacy profiles. The data presented here strongly suggest that the combination of selective inhibitors of BCL-XL plus MCL-1 and BCL-2 plus MCL-1 may be important new chemotherapeutic strategies in the management of cervical cancer.

## Acknowledgments

This work was supported by the Fundamental Research Grant Scheme, Ministry of Education Malaysia (203/PBIOLOGI/6711541), L’Oréal-UNESCO FWIS (304/PBIOLOGI/650853/L117) and Universiti Sains Malaysia RU grant (1001/PBIOLOGI/8012268).

