## Supplementary material for "Co-inhibition of BCL-XL and MCL-1 with BCL-2 selective inhibitors A1331852 and S63845 enhances cytotoxicity of cervical cancer cell lines": Table S1

**Table S1: $IC_{50}$ values of the cervical cancer cell lines to single agent treatment of A1331852, ABT-199 and S63845**

| Cell line | $IC_{50}$ (μM ± SEM) | | |
|---|---|---|---|
| | A1331852 | ABT-199 | S63845 |
| HeLa | 5.93 ± 0.8 | 14.63 ± 1.1 | 1.01 ± 0.16 |
| C33A | 20.47 ± 1.3 | 19.6 ± 1.9 | 7.72 ± 0.7 |
| SiHa | 24.06 ± 0.8 | 22.81 ± 0.8 | 11.41 ± 0.6 |
| CaSki | 1.61 ± 0.3 | 7.12 ± 0.6 | 5.72 ± 0.86 |
