## Supplementary material for "Co-inhibition of BCL-XL and MCL-1 with BCL-2 selective inhibitors A1331852 and S63845 enhances cytotoxicity of cervical cancer cell lines": Table S2

**Table S2: Sensitization of the cervical cancer cell lines to A1331852 by S63845.** The $IC_{50}$ values refer to the concentration of drug 1 **(bold)** that inhibits proliferation of 50% of the cells that were sensitized by the shown concentration of drug 2 *(italics)*. Fold sensitization: $IC_{50}$ drug 1/$IC_{50}$ drug 2.

| Cell line | **S63845 (µM)** | *$IC_{50}$ A1331852* (µM ± SEM) | Fold sensitization by S63845 |
|---|---|---|---|
| HeLa | **0** | 11.05 ± 1.95 | - |
| | **0.25** | *< 0.25** | 44 |
| | **0.5** | *< 0.25** | 44 |
| C33A | **0** | 24.66 ± 2.22 | - |
| | **0.5** | *< 0.25** | 99 |
| | **1** | *< 0.25** | 99 |
| SiHa | **0** | 25.13 ± 0.5 | - |
| | **0.5** | *< 0.25** | 100 |
| | **1** | *< 0.25** | 100 |
| CaSki | **0** | 1.61 ± 0.3 | - |
| | **0.5** | *< 0.25** | 6.4 |
| | **1** | *< 0.25** | 6.4 |

NOTE: *Where the parent $IC_{50}$ was not calculable, the lower bound was used.
