## Supplementary material for "Co-inhibition of BCL-XL and MCL-1 with BCL-2 selective inhibitors A1331852 and S63845 enhances cytotoxicity of cervical cancer cell lines": Table S3

**Table S3: The synergistic drug effects of A1331852 and S63845 in the cervical cancer cell lines.** All cervical cancer cell lines used were treated with increasing concentrations of A1331852 in the absence or presence of S63845.

| Cell Line | [S63845] (µM) | [A1331852] (µM) | CI | Drug combination interaction |
|---|---|---|---|---|
| HeLa | 0.25 | 0.25 | 0.042 | Very strong synergism |
| | | 0.5 | 0.039 | Very strong synergism |
| | | 1 | 0.037 | Very strong synergism |
| | | 2 | 0.05 | Very strong synergism |
| | | 4 | 0.064 | Very strong synergism |
| | 0.5 | 0.25 | 0.066 | Very strong synergism |
| | | 0.5 | 0.066 | Very strong synergism |
| | | 1 | 0.069 | Very strong synergism |
| | | 2 | 0.081 | Very strong synergism |
| | | 4 | 0.108 | Strong synergism |
| C33A | 0.5 | 0.25 | 0.013 | Very strong synergism |
| | | 0.5 | 0.012 | Very strong synergism |
| | | 1 | 0.015 | Very strong synergism |
| | | 2 | 0.012 | Very strong synergism |
| | | 4 | 0.012 | Very strong synergism |
| | 1 | 0.25 | 0.022 | Very strong synergism |
| | | 0.5 | 0.025 | Very strong synergism |

| | | | | |
|---|---|---|---|---|
| | | 1 | 0.022 | Very strong synergism |
| | | 2 | 0.023 | Very strong synergism |
| | | 4 | 0.022 | Very strong synergism |

| | | | | |
|---|---|---|---|---|
| | | 0.25 | 0.024 | Very strong synergism |
| | | 0.5 | 0.027 | Very strong synergism |
| | 0.5 | 1 | 0.037 | Very strong synergism |
| | | 2 | 0.057 | Very strong synergism |
| | | 4 | 0.099 | Very strong synergism |
| SiHa | | | | |
| | | 0.25 | 0.036 | Very strong synergism |
| | | 0.5 | 0.043 | Very strong synergism |
| | 1 | 1 | 0.049 | Very strong synergism |
| | | 2 | 0.07 | Very strong synergism |
| | | 4 | 0.11 | Strong Synergism |

| | | | | |
|---|---|---|---|---|
| CaSki | 0.5 | 0.25 | 0.072 | Very strong synergism |
| | | 0.5 | 0.107 | Strong synergism |
| | | 1 | 0.13 | Strong synergism |
| | | 2 | 0.23 | Strong synergism |
| | | 4 | 0.38 | Synergism |
| | 1 | 0.25 | 0.109 | Strong synergism |
| | | 0.5 | 0.115 | Strong synergism |
| | | 1 | 0.154 | Strong synergism |

| | | | |
|---|---|---|---|
| | 2 | 0.238 | Strong synergism |
| | 4 | 0.397 | Strong synergism |

NOTE: Combination index (CI) values were calculated using CompuSyn 1.0 (ComboSyn Inc., NJ, USA). [] indicates drug concentration; CI, combination index; values < 0.1 indicate very strong synergy and < 0.3 indicate strong synergy
