## Supplementary material for "Co-inhibition of BCL-XL and MCL-1 with BCL-2 selective inhibitors A1331852 and S63845 enhances cytotoxicity of cervical cancer cell lines": Table S4

**Table S4: Sensitization of the cervical cancer cell lines to ABT-199 by S63845.** The $IC_{50}$ values refer to the concentration of drug 1 **(bold)** that inhibits proliferation of 50% of the cells that were sensitized by the shown concentration of drug 2 *(italics)*. Fold sensitization: $IC_{50}$ drug 1/$IC_{50}$ drug 2.

| Cell line | **S63845 (µM)** | *$IC_{50}$ ABT-199* (µM ± SEM) | Fold sensitization by S63845 |
|---|---|---|---|
| HeLa | **0** | 22.05 ± 0.71 | - |
| | **0.25** | *6.34 ± 0.23* | 3 |
| | **0.5** | *1.65 ± 0.13* | 13 |
| C33A | **0** | 12.12 ± 1.39 | - |
| | **0.5** | *0.56 ± 0.12* | 22 |
| | **1** | *0.28 ± 0.03* | 43 |
| | **2** | *< 0.25** | 49 |
| SiHa | **0** | 27.84 ± 0.3 | - |
| | **0.5** | *12.5 ± 0.5* | 2.2 |
| | **1** | *10.26 ± 0.34* | 2.7 |
| | **2** | *8.88 ± 0.63* | 3.1 |
| CaSki | **0** | 7.12 ± 0.6 | - |
| | **0.5** | *1.24 ± 0.2* | 5.9 |
| | **1** | *0.52 ± 0.12* | 14.2 |
| | **2** | *0.47 ± 0.1* | 14.2 |

NOTE: *Where the parent $IC_{50}$ was not calculable, the lower bound was used.
