## Supplementary material for "Co-inhibition of BCL-XL and MCL-1 with BCL-2 selective inhibitors A1331852 and S63845 enhances cytotoxicity of cervical cancer cell lines": Table S5

**Table S5: The synergistic drug effects of ABT-199 and S63845 in the cervical cancer cell lines.** All cervical cancer cell lines used were treated with increasing concentrations of ABT-199 in the absence or presence of S63845.

| Cell Line | [S63845] (μM) | [ABT-199] (μM) | CI | Drug combination interaction |
|---|---|---|---|---|
| HeLa | 0.25 | 0.25 | ND | ND |
| | | 0.5 | ND | ND |
| | | 1 | ND | ND |
| | | 2 | 1.661 | Antagonism |
| | | 4 | 1.366 | Moderate antagonism |
| | 0.5 | 0.25 | 0.796 | Moderate synergism |
| | | 0.5 | 1.649 | Antagonism |
| | | 1 | 0.973 | Nearly additive |
| | | 2 | 0.428 | Synergism |
| | | 4 | 0.384 | Synergism |

| | | | | |
|---|---|---|---|---|
| C33A | 0.5 | 0.25 | 0.405 | Synergism |
| | | 0.5 | 0.426 | Synergism |
| | | 1 | 0.280 | Strong synergism |
| | | 2 | 0.214 | Strong synergism |
| | | 4 | 0.160 | Strong synergism |
| | 1 | 0.25 | 0.661 | Synergism |
| | | 0.5 | 0.526 | Synergism |
| | | 1 | 0.296 | Strong synergism |

| | | | | |
|---|---|---|---|---|
| | | 2 | 0.178 | Strong synergism |
| | | 4 | 0.112 | Strong synergism |
| | | 0.25 | 1.054 | Nearly additive |
| | | 0.5 | 0.848 | Moderate synergism |
| | 2 | 1 | 0.468 | Synergism |
| | | 2 | 0.254 | Strong synergism |
| | | 4 | 0.123 | Strong synergism |

| | | | | |
|---|---|---|---|---|
| | | 0.25 | ND | ND |
| | | 0.5 | ND | ND |
| | 0.5 | 1 | ND | ND |
| | | 2 | ND | ND |
| | | 4 | ND | ND |
| | | 0.25 | ND | ND |
| | | 0.5 | ND | ND |
| SiHa | 1 | 1 | ND | ND |
| | | 2 | ND | ND |
| | | 4 | ND | ND |
| | | 0.25 | ND | ND |
| | 2 | 0.5 | ND | ND |
| | | 1 | ND | ND |

| | | | | |
|---|---|---|---|---|
| | | 2 | ND | ND |
| | | 4 | ND | ND |
| | | 0.25 | 0.286 | Strong synergism |
| | | 0.5 | 0.311 | Synergism |
| | 0.5 | 1 | 0.314 | Synergism |
| | | 2 | 0.307 | Synergism |
| | | 4 | 0.179 | Strong synergism |
| | | 0.25 | 0.331 | Synergism |
| | | 0.5 | 0.267 | Strong synergism |
| CaSki | 1 | 1 | 0.264 | Strong synergism |
| | | 2 | 0.177 | Strong synergism |
| | | 4 | 0.148 | Strong synergism |
| | | 0.25 | 0.408 | Synergism |
| | | 0.5 | 0.455 | Synergism |
| | 2 | 1 | 0.288 | Strong synergism |
| | | 2 | 0.218 | Strong synergism |
| | | 4 | 0.221 | Strong synergism |

NOTE: Combination index (CI) values were calculated using CompuSyn 1.0 (ComboSyn Inc., NJ, USA). [] indicates drug concentration; CI, combination index; values < 0.1 indicate very strong synergy, < 0.3 indicate strong synergy, < 0.7 indicate synergy and < 0.85 indicate moderate synergy, < 0.9 indicate slight synergy, < 1.1 indicate nearly additive, < 1.2 indicate slight antagonism and < 1.45 indicate moderate antagonism; ND = not determined. CI values could not be determined for some drug concentrations as these concentrations were too low to cause an effect on cell proliferation. The words "antagonism", and "synergism" refer to the overall effect on cell proliferation and are not in any way meant to infer the properties of a classical pharmacological ligand that is an antagonist (in relation for example to a cell surface receptor and agonists/antagonists) pharmacological sense.
