## Supplementary figures and images for "Co-inhibition of BCL-XL and MCL-1 with BCL-2 selective inhibitors A1331852 and S63845 enhances cytotoxicity of cervical cancer cell lines"

### Fig S1

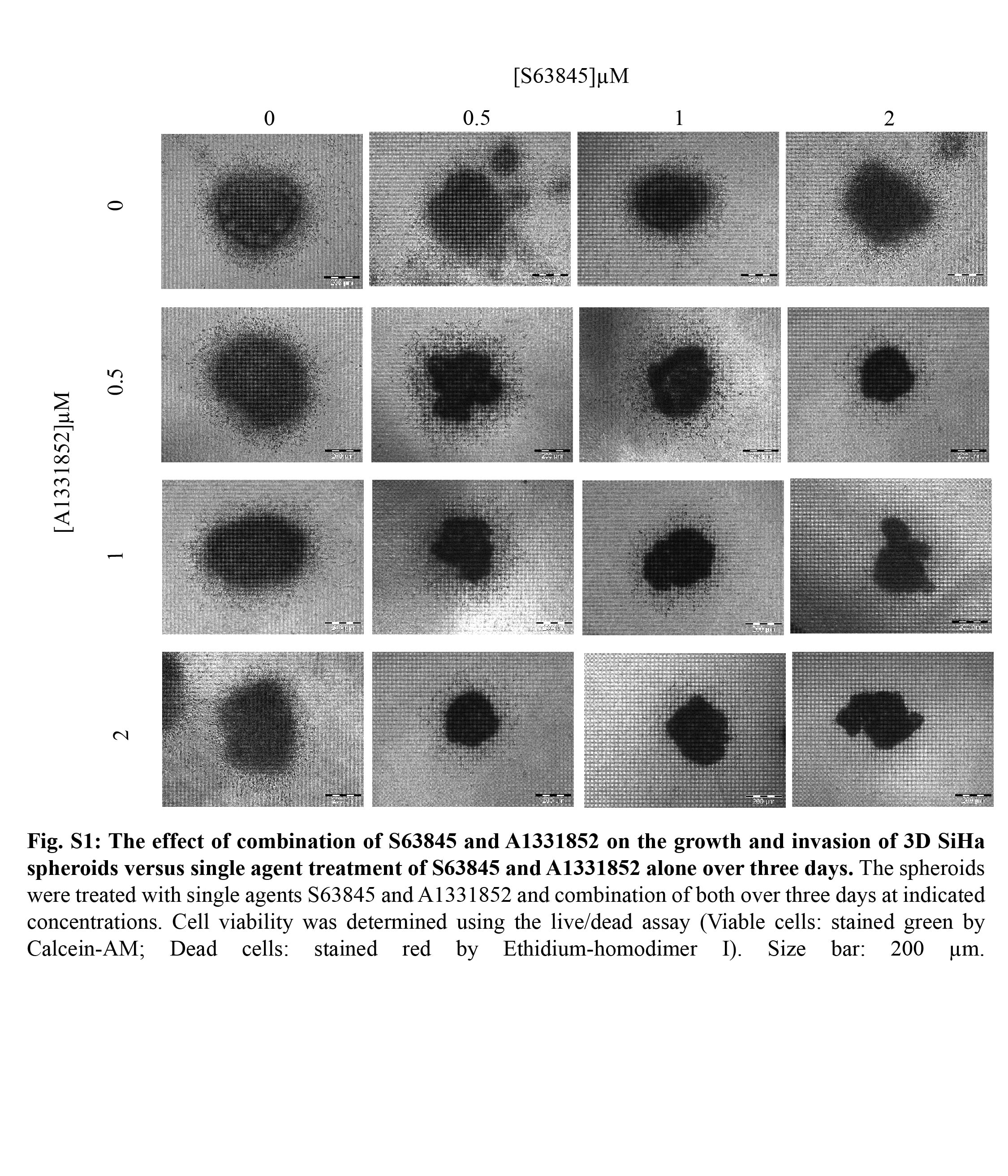

### Fig S2

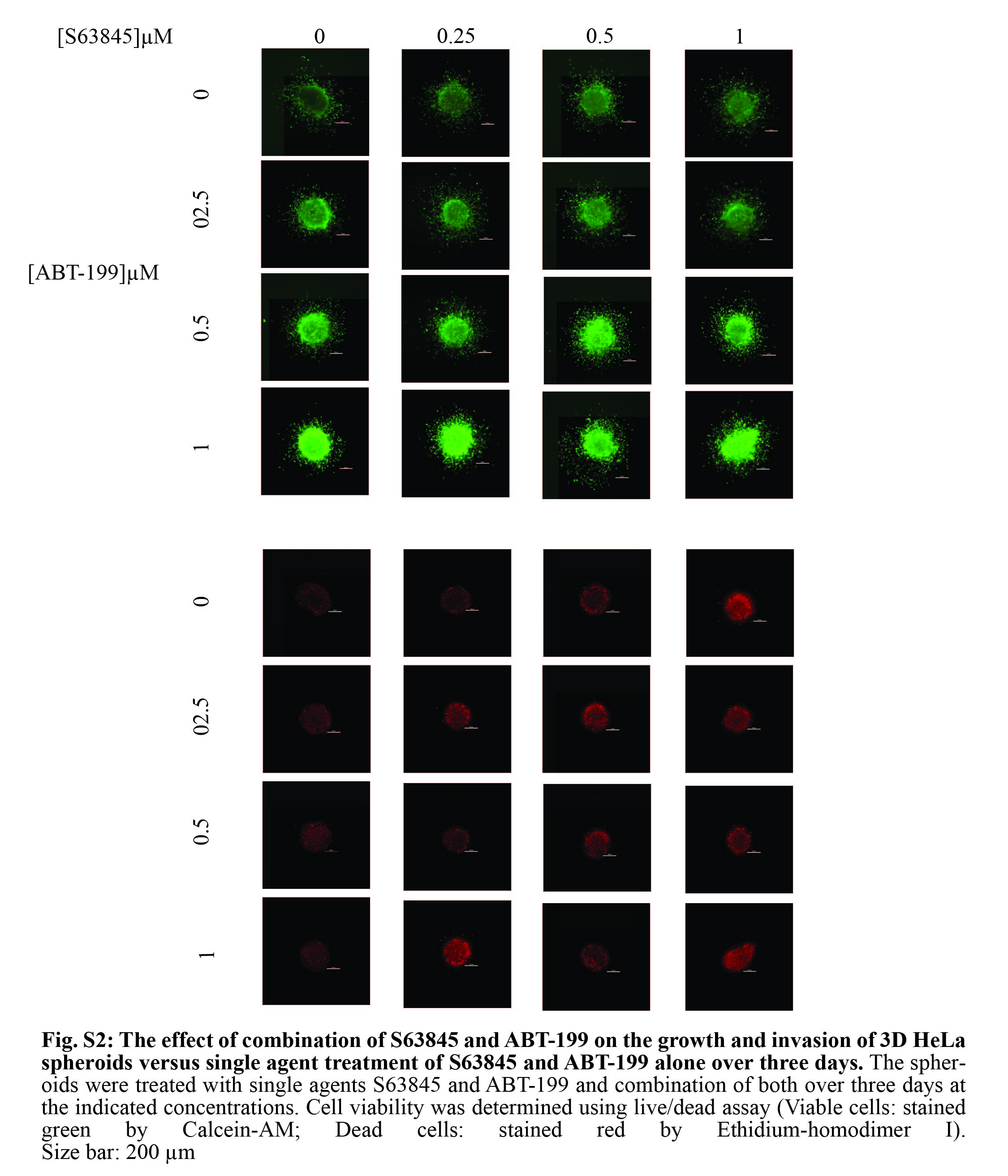
